# Health quality of seed potato and yield losses in Ecuador

**DOI:** 10.1101/108712

**Authors:** Israel Navarrete, Nancy Panchi, Peter Kromann, Gregory A. Forbes, Jorge L. Andrade-Piedra

## Abstract

Low potato productivity in Ecuador is partly attributed to the use of low quality seed tubers. However, seed health quality in Ecuador, and its interaction with altitude and yield has been poorly investigated. We surveyed 11 farmers’ fields in Ecuador in 2010 to determine incidence and severity of pathogens and pests affecting foliage and seed tubers, and to determine the influence of altitude and seed sources over seed health quality (pathogen and pest diversity in/on the seed tuber). Additionally, a field experiment was planted in CIP-Quito using assessed seed tubers collected from surveyed farmers’ fields, during 2010 and 2011, to determine yield responses to seed health quality. Results show that foliage was mainly affected by late blight and flea beetle damages while seed tubers were predominantly affected by black scurf, andean weevil damages, potato virus S and potato virus X. We found that seed health quality was similar among farmers’ seed sources, and detected that increase in altitude decreased seed-borne virus diversity. Only seed-borne pathogens and presence of mechanical damages were found to explain yield variation. Seed-borne pathogens affecting yield variation were black scurf on seed tubers, potato virus S, and potato yellow vein virus. However, these factors changed when regressions were performed per seed source or variety. The yield responses to seed health quality of each variety should be considered to fine-tune integrated pest management strategies.

## Introduction

Yield of the potato crop in Ecuador (mean = 7.9 t ha^-1^; standard deviation [sd]= 1.51; 1993 to 2014) is relatively low in comparison with neighbouring countries like Colombia (18.08 t ha^-1^) and Peru (14.7 t ha^-1^) (FAO 2016). This is caused, in part, by the use of a poor-quality seed (Thiele 1999), especially in terms of seed health (Fankhauser, 2000), as most farmers use seed tubers from informal sources, in which pathogens and pest accumulate over successive cycles of vegetative propagation, reducing yield and quality, in a process called *seed degeneration* (Thomas-Sharma et al. 2015). Viruses have been usually considered the main cause of degeneration, especially Potato Leafroll Virus (PLRV), Potato Virus A (PVA), Potato Virus S (PVS), Potato Virus X (PVX), and Potato Virus Y (PVY) (Salazar L.F. 2003; Scholthof et al. 2011). Other pathogens and pests that cause degeneration include *Rhizoctonia solani*, *Ralstonia solanacearum*, *Pectobacterium* spp., *Globodera* spp, (5) *Meloidogyne* spp., *Tecia solanivora*, among others (Thomas-Sharma et al. 2016).

The first report of seed degeneration in Ecuador was done more than a decade ago by Fankhauser (2000). The main results indicated that *R. solani*, *Streptomyces scabies* and *Premnotrypes vorax* were the most important pathogens and pests causing seed degeneration: incidence on tubers reached up to 78% for *R. solani*, 28% for *S. scabies*, and 33% for *P. vorax,* and yield reduction was between 17 and 30%. Results also showed low incidence on tubers of PLRV (<3%), PVY (<3%), PYVV (<2%), but not for PVX (up to 14%) and PVS (up to 96%). PYVV reduced yield in single plants from 21% to 41%, but not total yield. Yield losses caused by seed degeneration are highly variable and depend on the agro-ecological conditions, management and host genotype. Losses caused by viruses range from 10 to 90% depending on the virus. For instance, PVY can reduce the yield up to 90% and PVS up to 20% (Salazar L.F. 2003). However, in tropical highland areas where potatoes are grown above 2500 m.a.s.l, such as the Andes, viruses might not reduce the yield as much as in temperate regions, because of the negative correlation between altitude and temperature that decreases vector populations and virus dissemination (Sanchez de Luque et al. 1991; Bertschinger 1992; Fankhauser 2000). In the case of *R. solani* and *S.scabies*, yield losses have been estimated to be about 20 to 30% for *R. solani* (Banville 1989; Fankhauser 2000), and for *S. scabies* no data is available yet. In the case of pests, damages caused by the Andean potato weevil may reduce the price in about 20 to 50% (P. Oyarzún, Gallegos, et al. 2002), and the potato tuber moth complex (*Tecia solanivora*, *Phthorimaea operculella* and *Symmetrischema tangolias* [Dangles et al. 2009]) can cause yield losses of up to 40% and postharvest losses of up to 100% (Palacios et al. 1997). Despite the fact that we have information about the incidence of single pests, poor attention has been given to the multiple pathogens and pests affecting simultaneously the seed tubers.

Pathogen and pest incidence, severity and diversity on the tubers depends heavily on the seed source. In developing countries, most of the seed tubers comes from informal seed systems, including farmers’ own seed, neighbours or markets (Almekinders C. J. M. et al. 1994; Thiele 1999). In the case of Ecuador, the use of certified seed is increasing, from 3.3% of planted area in 2000 to 4.8% in 2015 (Instituto Nacional de Estadística y Censos 2016), but yields remain low (5.6 t ha^-1^ in 2000 and 7.3 t ha^-1^ in 2013) (FAO 2016). Perhaps, this low response of yield to the increased use of certified seed is because of seed quality, especially seed health, that is reduced in any part of the system. As consequence, farmers would not identify the benefits of using certified seed. Virus incidence in tubers coming from the formal system and the informal system in Peru and Ecuador suggest that this hypothesis is true (Bertschinger et al. 1990; Fankhauser 2000).

In this study, we report the results of one survey and one field experiment aiming at: (1) determining the incidence and severity of foliage and seed-borne pathogens and pests affecting the potato crop in Ecuador; (2) understanding how altitude and seed sources affect seed health quality; and (3) estimate the effect of seed health on yield. The implications of these results are discussed below.

## Materials and Methods

### Incidence and severity of seed borne pests

In 2010, eleven fields owned by small scale farmers of the Consortium of Small Potato Producers (CONPAPA in Spanish) were selected in three provinces of Ecuador (Table 1). The provinces were selected based on the number of hectares of potatoes cultivated, poverty levels, and presence of partner institutions. These were Bolivar, Chimborazo and Tungurahua. Fields were selected depending on the altitude (between 2751 and 3634 meters above the sea level, m.a.s.l) (Table 1), planted variety, and farmer’s eagerness to collaborate. Field area ranged between 220 and 1776 m^2^, and were planted with any of these six varieties: Chaucha Roja, Dolores, INIAP-Yana Shungo, INIAP-Fripapa, INIAP-Gabriela and Única. In ten fields (Table 1), incidence and severity of foliage diseases and insect damages were assessed in 100 randomly selected plants at flowering by using the keys proposed by Cruickshank et al. (1982) (number of plants per variety: Chaucha Roja = 200, Dolores = 100, INIAP-Fripapa = 200, INIAP-Gabriela = 200, Única = 200, INIAP-Yana Shungo = 100). Farmers were also asked to identify their seed sources. At harvest, farmers selected the best 400 tubers from each of the 11 fields and taken to the facilities of the International Potato Center in Quito (CIP-Quito, 3058 m.a.s.l, 0°22’ S, 78°33’ W). Tubers were washed, and incidence and severity of pests and diseases were visually assessed as described by James (1971) in each tuber. Physical and physiological issues, such as mechanical damages, misshapen tubers, and immature tubers were also considered. Nematodes were not considered in this survey. Number of tubers assessed per variety were: Chaucha Roja = 1017, Dolores = 400, INIAP-Fripapa = 800, INIAP-Gabriela = 800, Única = 855, INIAP-Yana Shungo = 400.

**Table 1.** Field location, potato varieties and seed sources in a survey conducted in the central highlands of Ecuador in 2010.

| Province | Altitude | Latitude | Longitude | Variety | Seed source<sup>1</sup> | Area (m<sup>2</sup>) |
| --- | --- | --- | --- | --- | --- | --- |
| **Bolívar** | 3596 | S 01° 32'52.4" | W 78° 55'14.5" | Dolores | Self-provided | 238 |
|  | 2812 | S 01° 38'38.8" | W 78° 58'29.1" | INIAP-Fripapa | Formal system (G3) | 803 |
|  | 2751 | S 01° 40'48.3" | W 78° 57'34.1" | Única | Self-provided | 1375 |
|  | 2793 | S 01° 40'45.0" | W 78° 57'26.8" | INIAP-Yana Shungo | Formal system (G4) | 220 |
| **Chimborazo** | 3576 | S 01° 36'31.6" | W 78° 48'21.8" | Chaucha Roja | Market | 888 |
|  | 3566 | S 01° 36'33.1" | W 78° 48'22.0" | Chaucha Roja | Market | 277 |
|  | 3455 | S 01° 36'28.2" | W 78° 46'49.3" | INIAP-Gabriela | Self-provided | 375 |
|  | 3601 | S 01° 38'28.9" | W 78° 47'48.4" | Chaucha Roja | Self-provided | 539 |
|  | 3527 | S 01° 34'25.1" | W 78° 46'37.1" | INIAP-Gabriela | Self-provided | 757 |
| **Tungurahua** | 3634 | S 01° 18'21.4" | W 78° 46'41.1" | INIAP-Fripapa | Formal system (G3) | 618 |
|  | 3035 | S 01° 06'26.7" | W 78° 32'00.2" | Única | Self-provided | 569 |

<sup>1</sup>The “G” notation refers to the number of successive plantings from in-vitro plants (Salazar 1995b). For the case of Ecuador, the seed G3 corresponds to the category “Registrada” and G4 to the category “Certificada”.

Tubers were then stored in diffuse light, covered with an anti-aphid net, and protected with insecticide (commercial name: Malathion® 50 PM; a.i, malathion, concentration: 500g/kg of product; dose: 5g of product/L) to prevent aphid infestation, until sprouting occurred naturally. A subsample of approximately 100 seed tubers was randomly selected out of the 400 seed tubers (selected per field) for virus diagnoses (number of tubers sampled per variety: Chaucha Roja = 404, Dolores = 103, INIAP-Fripapa = 240, INIAP-Gabriela = 237, Única = 286, INIAP-Yana Shungo = 90). To do this, one to 5 sprouts (2 to 3 cm long) of each tuber were collected depending on the availability and variety sprouting time (e.g., Chaucha Roja was collected at 29 days after farmers’ fields were harvested, while Única 110 days after harvest). Then, sprouts were disinfested (3% solution of Captan [i.a., Captan, dose; 1.5g/L idem]) and planted in trays in a net house at CIP-Quito, until having plants 8 to 10 cm tall. Plants were then transplanted into 3-kg pots and protected with an anti-aphid net until they were 15 to 20 cm tall. Leaf samples from top, middle and bottom of the plant were then taken and pooled together. Tubers used for virus diagnosis were put back into diffuse light storage until new sprouts appeared, and were used in the yield loss experiment described below. DAS-ELISA and NASH (Nucleic Acid Spot Hybridization) were performed for virus diagnosis using materials provided by the Virology Unit at CIP in Lima, Peru. DAS-ELISA was carried out according to the protocol proposed by CIP (2007) to detect the following viruses: Andean Potato Latent Virus (APLV), Andean Potato Mottle Virus (APMoV), PLRV, PVA, PVS, PVX, PVY. After completing DAS-ELISA, positive samples were determined by visual observation when samples turned yellow. Positive and negative controls were obtained from the National Agricultural Research Institute (acronym in Spanish INIAP) laboratory of Biotechnology and the BIOREBA company.

NASH was performed to detect Potato Yellow Vein Virus (PYVV) according to the protocols for Potato spindle tuber viroid proposed by CIP (1993a) and CIP (1993b). The samples were prepared in the facilities of CIP-Quito and then sent to the to the facilities of CIP in Lima for virus detection. To prepare the samples, briefly, nitrocellulose membranes were soaked in distilled water, washed twice with SSC 20X, and dried at ambient air. Then, leaf samples were macerated with two volumes of extraction buffer (49.4 ml of Formaldehyde (37%) + 50.6 ml of SSC 10X). These extracts were transferred to Eppendorf tubes and centrifugated at 12000 rpm per 5 minutes. After centrifugation, an aliquot of 3-4 μL of the supernatant was transferred to the membranes and dried at ambient air. Finally, membranes were baked at 80 °C for two hours.

Incidence was calculated as the percentage of plants with positive reaction in DAS-ELISA/nitrocellulose membranes in relation to the total number of plants.

### Yield losses caused by seed borne pests

A field experiment was carried out by planting a random subsample of 1360 seed tubers (experimental unit = one seed tuber) coming from the previous experiment (therefore pest incidence and severity and virus presence was known for each tuber) to estimate yield loss as a response of seed health, from October 2010 until December 2011 at CIP-Quito (average temperature 12.3°C and average accumulated rainfall per month 175 mm). Seed tubers were planted when sprouts were 1 to 2-cm long. Tubers from the same variety-field combination were planted together in 75-m^2^ plots, which were randomly allocated in a 2325-m^2^ plot. The fallow area between each 75-m^2^ plot was 2.7 m. Planting was done at 0.4 m between plants and 1.2 m between rows. Certified seed tubers (G4; [Salazar 1995]) of INIAP-Fripapa were planted between experimental units to avoid cross contamination. Plants were then labelled individually. Agronomic practices were done following local recommendations (P. Oyarzún, Gallegos, et al. 2002; P. Oyarzún, Chamorro, et al. 2002). Plants were harvested individually when reached full maturity (130 to 180 days after planting, depending on the variety) and yield was measured (g plant^-1^).

### Statistical analysis

Analysis of incidence and severity of foliage and seed borne pests were performed using descriptive statistics. Incidence data of seed borne pests were used to calculate the Shannon index, which provided an estimate of pathogen and pest diversity as a function of variety, seed source and altitude. It was calculated as described by Kosman (1996) and Perez et al. (2001) as follows *Hs* = − Σ_*j*_ (*p*_*j*_ x ln (*p*_*j*_), *j* = 1…*n*_*p*_, where *p*_*j*_ is the frequency of the *i*th seed borne pest and pathogen, and *n*_*p*_ is the number of seed borne pests and pathogens identified. An additional Shannon index was calculated only including viruses. Using the program “R” (version 3.3.2), a multiple linear regression was performed (command *lm*) to estimate the effect of seed health (pest and pathogen incidence and severity) of seed borne pests on yield loss. Assumptions of linearity were assessed by the default diagnostic plots of the program and the variance inflation factor (Car package, command *vif*) (Faraway 2016; Fox et al. 2016).

In order to deal with multicollinearity, a general regression model adjusting by the effects of the varieties was performed, and then the most meaningful parameters were selected. With these parameters, regressions were performed aggregating by seed source and variety. Raw data collected during the survey and field experiments (including weather data for the field experiment) was deposited in the CIP Dataverse Repository: http://dx.doi.org/10.21223/P3/XVAGXC (Navarrete et al. 2016)

## Results

During our survey, we identified three seed sources: (1) Self provided, (2) market, and (3) formal system (Table 1). The varieties Chaucha Roja, Dolores, INIAP-Gabriela and Única were self-provided by the farmers. Chaucha Roja from one farmer’s field was the only variety acquired in the market, and INIAP – Fripapa and INIAP- Yana Shungo (Table 1) were obtained from the formal system.

### Incidence and severity of foliage diseases, insect pests and seed borne pests

We identified 5 foliage diseases and 8 pests (Table 2). Diseases found were: black leg (*Pectobacterium* sp.), common rust of potato (*Puccinia pitteriana*), early blight (*Alternaria solani*), late blight (*Phytophthora infestans*), and stem canker (*Rhizoctonia solani*). Insect damages found were: Andean potato weevil (*Premnotrypes vorax*), aphids (several species involved such as *Macrosiphum* sp. or *Myzus* sp.), leafminer (*Liriomyza* spp.), potato flea beetle (*Epitrix* spp.), thrips (*Frankliniella tuberosi*), white grub (*Barotheus* sp), white fly (*Bemisia* sp. or *Trialeurodes vaporariorum*), and wireworm (*Agriotes* sp.). From this list, the most predominant diseases were late blight and common rust of potato. Average incidence of late blight was 65.9% and severity 15%. The lowest incidence was in variety INIAP-Yana Shungo (2%) and the highest was in Dolores (97%; Table 2). The second most prevalent disease was common rust of potato with an average incidence of 17% and severity of 1.3%. Varieties showing the lowest incidence were Chaucha roja and INIAP-Yana Shungo (0%); while the highest was found on the variety INIAP-Fripapa (49%;Table 2).

**Table 2.**
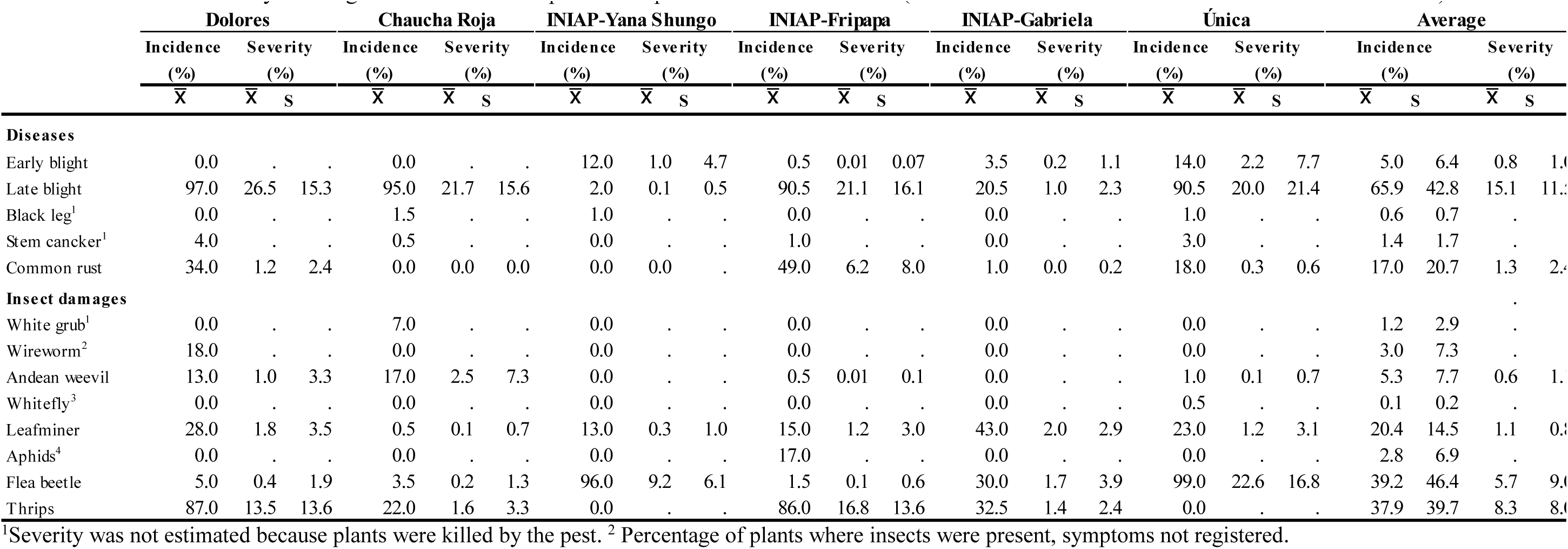
Incidence and severity of foliage diseases and insect pests in six potato varieties in Ecuador (for the number of observations see *Materials and Methods*)

|  | Dolores |  |  | Chaucha Roja |  |  | INIAP-Yana Shungo |  |  | INIAP-Fripapa |  |  | INIAP-Gabriela |  |  | Única |  |  | Average |  |  |  |
| --- | --- | --- | --- | --- | --- | --- | --- | --- | --- | --- | --- | --- | --- | --- | --- | --- | --- | --- | --- | --- | --- | --- |
|  | Incidence |  | Severity | Incidence |  | Severity | Incidence |  | Severity | Incidence |  | Severity | Incidence |  | Severity | Incidence |  | Severity | Incidence |  | Severity |  |
|  | (%) |  | (%) | (%) |  | (%) | (%) |  | (%) | (%) |  | (%) | (%) |  | (%) | (%) |  | (%) | (%) |  | (%) |  |
| | $\bar{X}$ | $\bar{X}$ | s | $\bar{X}$ | $\bar{X}$ | s | $\bar{X}$ | $\bar{X}$ | s | $\bar{X}$ | $\bar{X}$ | s | $\bar{X}$ | $\bar{X}$ | s | $\bar{X}$ | $\bar{X}$ | s | $\bar{X}$ | s | $\bar{X}$ | s |
| **Diseases** |  |  |  |  |  |  |  |  |  |  |  |  |  |  |  |  |  |  |  |  |  |  |
| Early blight | 0.0 | . | . | 0.0 | . | . | 12.0 | 1.0 | 4.7 | 0.5 | 0.01 | 0.07 | 3.5 | 0.2 | 1.1 | 14.0 | 2.2 | 7.7 | 5.0 | 6.4 | 0.8 | 1.0 |
| Late blight | 97.0 | 26.5 | 15.3 | 95.0 | 21.7 | 15.6 | 2.0 | 0.1 | 0.5 | 90.5 | 21.1 | 16.1 | 20.5 | 1.0 | 2.3 | 90.5 | 20.0 | 21.4 | 65.9 | 42.8 | 15.1 | 11.5 |
| Black leg<sup>1</sup> | 0.0 | . | . | 1.5 | . | . | 1.0 | . | . | 0.0 | . | . | 0.0 | . | . | 1.0 | . | . | 0.6 | 0.7 | . | . |
| Stem canker<sup>1</sup> | 4.0 | . | . | 0.5 | . | . | 0.0 | . | . | 1.0 | . | . | 0.0 | . | . | 3.0 | . | . | 1.4 | 1.7 | . | . |
| Common rust | 34.0 | 1.2 | 2.4 | 0.0 | 0.0 | 0.0 | 0.0 | 0.0 | . | 49.0 | 6.2 | 8.0 | 1.0 | 0.0 | 0.2 | 18.0 | 0.3 | 0.6 | 17.0 | 20.7 | 1.3 | 2.4 |
| **Insect damages** |  |  |  |  |  |  |  |  |  |  |  |  |  |  |  |  |  |  |  |  |  |  |
| White grub<sup>1</sup> | 0.0 | . | . | 7.0 | . | . | 0.0 | . | . | 0.0 | . | . | 0.0 | . | . | 0.0 | . | . | 1.2 | 2.9 | . | . |
| Wireworm<sup>2</sup> | 18.0 | . | . | 0.0 | . | . | 0.0 | . | . | 0.0 | . | . | 0.0 | . | . | 0.0 | . | . | 3.0 | 7.3 | . | . |
| Andean weevil | 13.0 | 1.0 | 3.3 | 17.0 | 2.5 | 7.3 | 0.0 | . | . | 0.5 | 0.01 | 0.1 | 0.0 | . | . | 1.0 | 0.1 | 0.7 | 5.3 | 7.7 | 0.6 | 1.0 |
| Whitefly<sup>3</sup> | 0.0 | . | . | 0.0 | . | . | 0.0 | . | . | 0.0 | . | . | 0.0 | . | . | 0.5 | . | . | 0.1 | 0.2 | . | . |
| Leafminer | 28.0 | 1.8 | 3.5 | 0.5 | 0.1 | 0.7 | 13.0 | 0.3 | 1.0 | 15.0 | 1.2 | 3.0 | 43.0 | 2.0 | 2.9 | 23.0 | 1.2 | 3.1 | 20.4 | 14.5 | 1.1 | 0.8 |
| Aphids<sup>4</sup> | 0.0 | . | . | 0.0 | . | . | 0.0 | . | . | 17.0 | . | . | 0.0 | . | . | 0.0 | . | . | 2.8 | 6.9 | . | . |
| Flea beetle | 5.0 | 0.4 | 1.9 | 3.5 | 0.2 | 1.3 | 96.0 | 9.2 | 6.1 | 1.5 | 0.1 | 0.6 | 30.0 | 1.7 | 3.9 | 99.0 | 22.6 | 16.8 | 39.2 | 46.4 | 5.7 | 9.0 |
| Thrips | 87.0 | 13.5 | 13.6 | 22.0 | 1.6 | 3.3 | 0.0 | . | . | 86.0 | 16.8 | 13.6 | 32.5 | 1.4 | 2.4 | 0.0 | . | . | 37.9 | 39.7 | 8.3 | 8.0 |

<sup>1</sup>Severity was not estimated because plants were killed by the pest. <sup>2</sup>Percentage of plants where insects were present, symptoms not registered.

Potato flea beetle and thrips were predominantly found during the survey (Table 2). The average incidence of the potato flea beetle was 39.2% and the severity 5.7% with the lowest incidence found on INIAP-Fripapa (1.5%) and the highest on Única (99%). Average incidence of thrips was 37.9%. We did not find thrips on the plants of INIAP-Yana Shungo, but these were found on 87% of plants of the variety Dolores. We found 19 seed borne pests and three physical and physiological blemishes (Table 3). Diseases found on the seed tubers were: black leg (*Pectobacterium* sp.), black scurf (*Rhizoctonia solani*), fusarium dry rot (*Fusarium* sp.), silver scurf (*Helminthosporium* sp.), powdery scab (*Spongospora subterranea*), and cracking, i.e., term used to describe symptoms caused by *R. solani* (elephant hide), *Streptomyces* spp. and other cracking like symptoms. Insect damage was caused by the Andean potato weevil (*Premnotrypes vorax*), potato flea beetle (*Epitrix* sp.), potato tuber moth complex (*Tecia solanivora*, *Symmetrischema tangolias*, or *Phthorimaea operculella*), white grub (*Barotheus* spp.), and wireworm (*Agriotes* spp.). The most prevalent diseases were black scurf and cracking (Table 3). Average incidence of black scurf was 80.5% and severity was 3.2%. The lowest incidence of black scurf was in variety INIAP-Yana Shungo (42%) and the highest was in Única (95.8%). Cracking was the second most important problem affecting seed health quality. Average incidence was 18.3% and average severity was 0.6%. Chaucha Roja was the least affected variety (1.5% incidence) while INIAP-Gabriela was the most affected (37.3%).

**Table 3.** Incidence and severity of seed borne diseases, insect pests, and physical and physiological blemishes in six potato varieties in Ecuador (for the number of observations see *Materials and Methods*).

|  | Dolores |  |  | Chaucha Roja |  |  | INIAP-Yana Shungo |  |  | INIAP-Fripapa |  |  | INIAP-Gabriela |  |  | Única |  |  | Average |  |  |  |
| --- | --- | --- | --- | --- | --- | --- | --- | --- | --- | --- | --- | --- | --- | --- | --- | --- | --- | --- | --- | --- | --- | --- |
|  | Incidence |  | Severity | Incidence |  | Severity | Incidence |  | Severity | Incidence |  | Severity | Incidence |  | Severity | Incidence |  | Severity | Incidence |  | Severity |  |
|  | (%) |  | (%) | (%) |  | (%) | (%) |  | (%) | (%) |  | (%) | (%) |  | (%) | (%) |  | (%) | (%) |  | (%) |  |
| | $\bar{X}$ | $\bar{X}$ | s | $\bar{X}$ | $\bar{X}$ | s | $\bar{X}$ | $\bar{X}$ | s | $\bar{X}$ | $\bar{X}$ | s | $\bar{X}$ | $\bar{X}$ | s | $\bar{X}$ | $\bar{X}$ | s | $\bar{X}$ | s | $\bar{X}$ | s |
| Diseases |  |  |  |  |  |  |  |  |  |  |  |  |  |  |  |  |  |  |  |  |  |  |
| Cracking | 14.5 | 0.3 | 1.0 | 1.5 | 0.0 | 0.4 | 8.0 | 0.1 | 0.3 | 33.3 | 1.1 | 3.6 | 37.3 | 1.6 | 5.0 | 15.5 | 0.6 | 3.7 | 18.3 | 14.1 | 0.6 | 0.6 |
| Fusarium dry rot | 3.0 | 0.5 | 5.7 | 35.8 | 3.0 | 7.3 | 3.0 | 0.2 | 1.4 | 6.8 | 0.3 | 1.8 | 0.3 | 0.0 | 0.3 | 2.8 | 0.6 | 6.6 | 8.6 | 13.5 | 0.8 | 1.1 |
| Silver scurf | 0.0 | 0.0 | . | 0.0 | 0.0 | . | 0.0 | 0.0 | . | 4.8 | 0.3 | 1.6 | 0.0 | 0.0 | . | 39.5 | 5.3 | 12.6 | 7.4 | 15.9 | 0.9 | 2.1 |
| Black leg | 0.0 | 0.0 | . | 1.0 | 0.8 | 8.0 | 2.0 | 0.5 | 4.1 | 0.0 | 0.0 | . | 0.0 | 0.0 | . | 0.3 | 0.1 | 1.3 | 0.5 | 0.8 | 0.2 | 0.3 |
| Black scurf | 88.5 | 5.1 | 6.2 | 83.3 | 2.6 | 3.3 | 42.0 | 0.4 | 0.6 | 85.0 | 2.4 | 6.8 | 88.3 | 3.8 | 7.4 | 95.8 | 4.6 | 6.5 | 80.5 | 19.3 | 3.2 | 1.7 |
| Powdery scab | 0.0 | 0.0 | . | 0.0 | 0.0 | . | 0.0 | 0.0 | . | 0.3 | 0.0 | 0.5 | 0.5 | 0.0 | 0.1 | 0.0 | 0.0 | . | 0.1 | 0.2 | 0.0 | 0.0 |
| Virus |  |  |  |  |  |  |  |  |  |  |  |  |  |  |  |  |  |  |  |  |  |  |
| APLV | 0.0 | . | . | 0.0 | . | . | 19.5 | . | . | 0.0 | . | . | 9.0 | . | . | 3.4 | . | . | 5.3 | 7.8 | . | . |
| APMoV | 0.0 | . | . | 0.0 | . | . | 0.0 | . | . | 0.0 | . | . | 6.0 | . | . | 0.0 | . | . | 1.0 | 2.4 | . | . |
| PLRV | 0.0 | . | . | 0.0 | . | . | 2.4 | . | . | 1.0 | . | . | 1.0 | . | . | 5.6 | . | . | 1.7 | 2.1 | . | . |
| PVA | 0.0 | . | . | 0.0 | . | . | 0.0 | . | . | 0.0 | . | . | 5.0 | . | . | 0.0 | . | . | 0.8 | 2.0 | . | . |
| PVS | 100.0 | . | . | 18.0 | . | . | 85.4 | . | . | 0.0 | . | . | 4.0 | . | . | 69.7 | . | . | 46.2 | 44.0 | . | . |
| PVX | 88.0 | . | . | 100.0 | . | . | 0.0 | . | . | 2.0 | . | . | 30.0 | . | . | 42.7 | . | . | 43.8 | 42.4 | . | . |
| PVY | 2.0 | . | . | 0.0 | . | . | 2.4 | . | . | 2.0 | . | . | 9.0 | . | . | 1.1 | . | . | 2.8 | 3.2 | . | . |
| PYVV | 0.0 | . | . | 0.0 | . | . | 2.4 | . | . | 2.0 | . | . | 11.0 | . | . | 1.1 | . | . | 2.8 | 4.2 | . | . |
| Insects damages |  |  |  |  |  |  |  |  |  |  |  |  |  |  |  |  |  |  |  |  |  |  |
| White grub | 10.5 | 0.4 | 1.8 | 5.0 | 0.2 | 1.7 | 16.5 | 1.8 | 7.4 | 1.0 | 0.0 | 0.6 | 6.8 | 0.3 | 1.5 | 10.0 | 0.9 | 4.2 | 8.3 | 5.3 | 0.6 | 0.7 |
| Wireworm | 11.5 | 0.2 | 0.7 | 1.3 | 0.0 | 0.3 | 0.0 | 0.0 | . | 8.5 | 0.1 | 0.7 | 15.8 | 0.3 | 1.0 | 0.0 | 0.0 | 0.0 | 6.2 | 6.7 | 0.1 | 0.1 |
| Andean weevil | 82.5 | 14.4 | 18.8 | 61.8 | 4.2 | 9.1 | 33.5 | 1.7 | 4.5 | 45.3 | 3.4 | 7.8 | 53.8 | 4.2 | 7.8 | 1.5 | 0.1 | 0.4 | 46.4 | 27.5 | 4.7 | 5.0 |
| Tuber moth complex | 0.0 | 0.0 | . | 2.8 | 0.1 | 0.5 | 0.0 | 0.0 | . | 0.0 | 0.0 | 0.0 | 0.0 | 0.0 | . | 25.3 | 1.0 | 3.8 | 4.7 | 10.1 | 0.2 | 0.4 |
| Flea beetle | 0.0 | 0.0 | . | 0.0 | 0.0 | . | 0.0 | 0.0 | . | 0.0 | 0.0 | . | 1.0 | 0.2 | 2.6 | 8.3 | 0.2 | 1.0 | 1.5 | 3.3 | 0.1 | 0.1 |
| Physical and physiological issues |  |  |  |  |  |  |  |  |  |  |  |  |  |  |  |  |  |  |  |  |  |  |
| Mechanical damage | 2.5 | . | . | 10.0 | . | . | 10.5 | . | . | 14.5 | . | . | 13.5 | . | . | 20.8 | . | . | 12.0 | 6.0 | . | . |
| Misshappen tubers | 0.5 | . | . | 1.5 | . | . | 12.5 | . | . | 2.3 | . | . | 0.0 | . | . | 1.5 | . | . | 3.0 | 4.7 | . | . |
| Inmature tubers | 35.0 | . | . | 51.5 | . | . | 99.5 | . | . | 0.8 | . | . | 63.0 | . | . | 33.3 | . | . | 47.2 | 33.2 | . | . |

Andean weevil and white grub damages were also important (Table 3). Average incidence of the damage of Andean weevil was 46.4% with a severity of 4.7%. The least affected variety was Única (1.5% incidence) and the most affected was Dolores (82.5%). Damage caused by white grubs was found in 8.3% of tubers with a severity of 0.6% (Table 3). The least affected variety was INIAP-Fripapa (1% of incidence) and the most affected variety was INIAP-Yana Shungo (16.5%). Tubers with symptoms of tuber blight (*P. infestans*) were not found in this study.

Virus diagnosis revealed that 38.0% of the tubers sampled were free of virus, 45.1% were infected with one virus, 16.3% were infected with two viruses, and 0.01% were infected with three viruses. Virus infection with more than three viruses was not found on the sample. Viruses with the highest incidence were PVS (46.2%) and PVX (43.8%) (Table 3). PVS was not found on INIAP-Fripapa, but in all the tubers of Dolores (Table 3). Similarly, PVX was not found on INIAP-Yana Shungo, but it was found in all the tubers of Chaucha Roja. Incidences of PVA and APMoV were low (0.8 and 1%, respectively; Table 3). These two viruses were found only in the variety INIAP-Gabriela in 5% and 6% of the tubers, respectively. PVX and PVS were present simultaneously in 11% of the tubers. Other simultaneous viral infections were low (incidence < 2%), these were: APLV+PLRV=0.7%, APLV+PVS=1.7%, APLV+PVX=1%, APLV+PVY=0.4%, APMoV+PVA=0.8%, APMoV+PVS=0.2%, APMoV+PVX=0.1%, APMoV+PYVV=0.1%, PLRV+PVX=0.8%, PVA+PVX=0.2%, PVS+PVY=0.4%, PVS+PYVV=0.5%, PVX+PVY=0.4%, PVX+PYVV=0.5, PYV+PYVV=0.2%. PVX+PVY+PVS=0.001%, PVX+PVS+PYVV<0.001%, PVX+PVS+APLV<0.001%, PVX+PLRV+APLV<0.001%, PVX+APMoV+PVA<0.001%, PVY+PVS+APLV=0.001%.

Three physical and physiological blemishes were found during this survey: mechanical damages, misshapen tubers, and inmature tubers. Interestingly, almost half of the tubers sampled were found to be inmature (47.2%). The lowest incidence of inmature tubers was found on the variety INIAP-Fripapa (0.8%), while the highest was found on the variety INIAP-Yana Shungo (99.5%). A small portion of misshapen tubers were found during this sampling (3%, Table 3). All the tubers of the variety INIAP-Gabriela had the variety’s tuber shape at harvest, while 20.8% of the tubers of the variety Única were misshapen.

### Seed borne pest and pathogen diversity

Pest and pathogen diversity on the seed tubers was assessed using the Shannon diversity index where a larger index indicates a larger diversity. Taking into account all pests and diseases, the Shannon diversity indexes were the following: Dolores: 1.12 (*n* = 103 tubers); INIAP-Fripapa: 1.62 (*n* = 240); Chaucha Roja: 1.94 (*n* = 404); INIAP-Yana Shungo: 2.18 (*n* = 90); Única: 2.39 (*n* = 286); and INIAP-Gabriela: 2.71 (*n* = 237). The Shannon diversity index for all the pests did not show any relationship with the altitude. However, we found a negative correlation with altitude when the Shannon diversity index was calculated for viruses only (Figure 1A). The coefficient of determination (R^2^) indicated that the altitude explained 6% of the variability of the Shannon diversity index when including all varieties (Figure 1A, dashed line), and 22% when excluding INIAP-Gabriela (Figure 1A, full line). Considering all the pests and pathogens, we found no differences in the Shannon diversity index among tubers harvested from different seed sources (*p* > 0.05) (Figure 1B): tubers produced with seed from the formal system had a Shannon diversity index of 1.7 (sd = 0.5; *n* = 330 tubers); tubers coming from seed from the market: 1.67 (sd = 0.6; *n* = 360); and tubers coming from seed which was self-provided by the farmer: 1.82 (sd = 0.6; *n* = 670).

**Figure 1.**
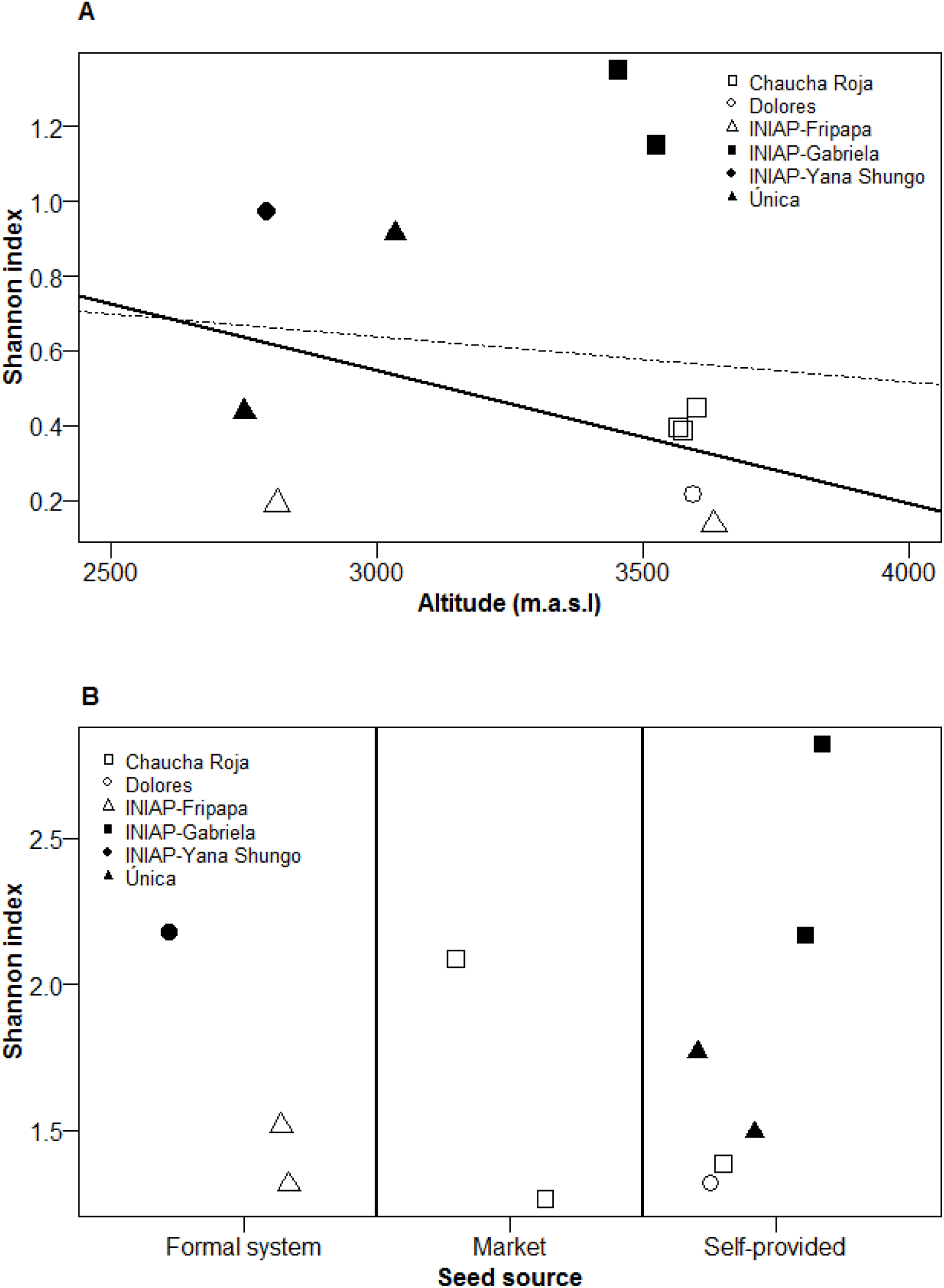
Seed borne pest diversity on seed tubers surveyed in three provinces in Ecuador. A. Effect of the altitude on virus diversity: dashed line represents a model including all varieties (R^2^ = 6%), and full line a model excluding INIAP-Gabriela (R^2^ = 22%). B. Pest diversity on tubers according to the three seed sources.

### Yield losses caused by seed borne pests

Average yield during the field experiment was 1010.5 g plant^-1^ (sd= 785.8) and, as expected, there were large differences among varieties: Chaucha Roja: 437.1 g plant^-1^ (sd = 298.4); Dolores: 595.8 g plant^-1^ (sd = 252.7); INIAP-Fripapa 1150.1 g plant^-1^ (sd = 417.9); INIAP-Gabriela: 2262.8 g plant^-1^ (sd = 717.1); INIAP-Yana Shungo: 883.8 g plant^-1^ (sd = 515.8); and Única: 734.9 g plant^-1^ (sd= 454.7).

Before describing the models obtained in this section, it is worth to note that the factor of varieties was considered important driving yield variation. It was able to explain 65% of the variability in yield, but it was excluded from the analysis due to problems of multicollinearity.

The multiple linear regression identified (*n* = 1103 observations) four significant parameters interacting with the varieties studied: presence or absence of PVS, PYVV and mechanical damage, and severity (%) of black scurf on the seed tuber (Table 4). Multiple regressions aggregated depending on the seed source were different. The multiple linear regression for the seed that was self-provided by the farmer had an R^2^ (coefficient of determination) of 40%, and predicted and average yield of 1739.2. It detected the presence and absence of PVS and PYVV as important factors explaining yield variability. PVS had a negative influence over the yield explaining 38% of the variability while the presence or absence of PYVV has a positive influence on yield explaining 2% of the variability (Table 4). The model estimated for the varieties acquired in the market showed a lower R^2^ of 7% with an average yield of 375.3 g plant^-1^. It identified the severity of black scurf as the only important factor affecting positively yield (Table 4). The regression for the seed that was acquired from the formal system had an R^2^ of 7% and estimated an average yield of 1181.6 g plant^-1^. The model identified two significant parameters that had a negative influence on yield, presence and absence of PVS and mechanical damage (Table 4).

**Table 4.** Main seed borne pests affecting yield variation. Quito, Pichincha.2012

| Parameter<sup>1,2</sup> | Observations | Coefficient<br>(g plant<sup>-1</sup>) | Partial R-squared<br>(%) |
| --- | --- | --- | --- |
| **Models per seed source** |  |  |  |
| **Self-provided** | 539 |  |  |
| Intercept |  | 1739.2 |  |
| PVS |  | -1180.0 | 38 |
| PYVV |  | 815.2 | 2 |
| Total R-squared |  |  | 40 |
| **Market** | 267 |  |  |
| Intercept |  | 375.3 |  |
| Black scurf |  | 24.0 | 7 |
| Total R-squared |  |  | 7 |
| **Formal system** | 297 |  |  |
| Intercept |  | 1181.6 |  |
| PVS |  | -280.7 | 5 |
| Mechanical damage |  | -184.4 | 2 |
| Total R-squared |  |  | 7 |
| **Models per variety** |  |  |  |
| **Chaucha Roja** | 292 |  |  |
| Intercept |  | 388.6 |  |
| PVS |  | -111.5 | 1 |
| Black scurf |  | 23.8 | 7 |
| Total R-squared |  |  | 8 |
| **INIAP-Fripapa** | 221 |  |  |
| Intercept |  | 1220.6 |  |
| Mechanical damage |  | -299.6 | 9 |
| Total R-squared |  |  | 9 |
| **INIAP-Gabriela** | 197 |  |  |
| Intercept |  | 2178.9 |  |
| PVS |  | -944.1 | 3 |
| Black scurf |  | 18.8 | 2 |
| PYVV |  | 526.1 | 4 |
| Total R-squared |  |  | 9 |
| **INIAP- Yana Shungo** | 76 |  |  |
| Intercept |  | 835.8 |  |
| Mechanical damage |  | 730.0 | 11 |
| Total R-squared |  |  | 11 |
| **Única** | 223 |  |  |
| Intercept |  | 925.8 |  |
| PVS |  | -251.4 | 8 |
| Black scurf |  | 7.2 | 3 |
| Total R-squared |  |  | 11 |

<sup>1</sup> Parameters for multiple linear regressions were selected when *p-value* was lower 0.05. *P- values* ranged from < 0.001 to 0.03.
<sup>2</sup> PVS, PYVV, and mechanical damage were assessed based on their presence and absence; while black scurf was based on the percentage of severity on the seed tuber.

Regressions were different among varieties which R^2^ ranged from 8 to 11% (Table 4). For Chaucha Roja, the model (R^2^ = 8%) found the presence and absence of PVS and severity of black scurf as significant predictors affecting yield. PVS had a negative influence on yield while severity of black scurf had a positive influence on yield. For INIAP-Fripapa (R^2^ = 9%), the regression identified only the presence or absence of mechanical damage as significant parameters for yield variation. For INIAP-Gabriela (R^2^ = 14%), three predictors were detected: presence or absence of PVS, presence or absence of PYVV, and severity of black scurf. PVS had a negative effect on yield variation while black scurf and PYVV had a positive effect on yield variation. For variety INIAP-Yana Shungo (R^2^ = 11%), the model identifies the presence or absence of mechanical damage as the only predictor of yield variability. For Unica (R^2^ = 11%), the regression detected two predictors: presence or absence of PVS and severity (%) of black scurf. PVS had a negative effect on yield variation while black scurf had a positive effect on yield variation. The model for variety Dolores was not estimated since the contribution of the parameters to the variation in yield was not significant.

## Discussion

Main findings of this study were the following: (1) in the foliage, the most important pests were late blight and potato flea beetle (Table 2), while in the tubers the most important were black scurf and Andean potato weevil; viruses with the highest incidences were PVS and PVX; (2) altitude had a negative effect on virus diversity and explained between 6 and 22% of its variability; pest diversity was similar in tubers harvested from different seed sources; (3) models estimating yield responses were different when aggregated by seed source or varieties. Due to the fact that the survey and the field experiment were performed in a single year (2010-2011) omitting replicates, conclusions from this study have to be taken cautiously.

### Incidence and severity of seed borne pests

The most important foliage disease found was late blight. This disease was present in 65.9% of the plants assessed (Table 2), indicating that it is still one of the main biotic factors limiting the productivity in the country as suggested by Hijmans et al. (2000) and P. Oyarzún et al. (2001). Damages of potato flea beetle follow late blight in importance with an incidence 39.2%. Although, we were aware about the importance about this pest, it is the first time that data about damage of potato flea beetle is reported in Ecuador.

Black scurf and Andean potato weevil showed the highest incidences on tubers. Black scurf was present in the 80.5% of the tubers sampled, however, low severities were detected (mean = 3.2%, sd = 1.7, Table 3). Fankhauser (2000) reported similar results about incidence of black scurf when sampling tubers of the varieties INIAP-Gabriela and INIAP-Esperanza in the province Chimborazo (78% incidence). Incidence of the Andean potato weevil was also high (46.4%) (Table 3), but approximately 50% less than previously reported by Fankhauser (2000) (88% incidence). These results may indicate that management strategies to control *R. solani* proposed by local organizations are not working, or are not being adopted by farmers since the incidence of black scurf remain stable from 1998 to 2010, when Fankhauser (2000) and our group did the surveys. Our experience suggest a poor adoption of the management strategies by farmers, and points out the need of strengthening local extension services (Parsa et al. 2012). In contrast, incidence of Andean potato weevil has dropped nearly 50%, from 88% in 1998 to 43.9% in 2010, suggesting better adoption of integrated management practices, more efficient insecticides, or other causes (e.g., an unforeseen effect of climate change on the population of *Premnotrypes vorax)*.

Cracking symptoms were found on the 18.3% of the tubers sampled (Table 3). This high incidence reported might be the result of the use of “cracking” as a generic term to refer to symptoms that could be produced either by *R. solani, Streptomyces* spp., or other causes, and suggest the need for a specific study to define the etiology of cracking.

Our results confirm that the presence of tuber blight in the country is limited. A previous research found similar incidences of tuber blight on farmer fields in the provinces of Bolivar, Cañar, Carchi, Chimborazo, Pichincha and Tungurahua (P. J. Oyarzún et al. 2005). Probably, the low incidence of tuber blight found is associated with the soil microbiological and physical-chemical characteristics affecting the development of the disease (Carla Garzón and Forbes 1999; Villamarin et al. 2011). In spite of it, special attention should be given to tuber blight since there is evidence of pre-emergence infection caused by *P. infestans* in Ecuador (Kromann et al. 2008).

Viruses have been considered the main factors to reduce seed quality. Our results showed that in the provinces of Bolivar, Chimborazo and Tungurahua of Ecuador, the mild viruses PVS and PVX were the most prevalent viral problems, found in 46.2% and 43.8% of the tubers sampled, respectively (Table 3). Surveys performed in Peru during 1985-1987 and in Ecuador in 1998 also found that both of these viruses were predominant in farmers’ fields (Bertschinger et al. 1990; Fankhauser 2000). On the contrary, recent research points out to a high incidence of PVY in the varieties INIAP-Fripapa and Superchola (Gomez et al. 2015), but such contrast might be influenced by the seed source. This variation in results stresses out the importance of understanding ecological conditions, on-farm management practices, seed sources and planted variety to determine viral incidence.

### Seed borne pest and pathogen diversity

We found that altitude has a negative effect on virus diversity. Our results support the findings of Sanchez de Luque et al. (1991) and Fankhauser (2000) where they report higher incidences of PVX, PVS, PVY, PLRV and PYVV at lower altitudes than at higher altitudes. Our results confirm that there are beneficial effects of the traditional on-farm practice of using seed tubers produced at high altitudes as an strategy to improve seed health (Thiele 1999). However, specific experiments should be considered to confirm these findings, and determine the effect of the seed tubers produced at higher altitudes when planted at lower altitudes or other plots on the productivity and seed health quality.

Our results show that, at the time the survey was done, seed health was similar among seed sources (Figure 1B). Similar observation were done by Bertschinger et al. (1990) when comparing viral incidence among different seed sources. However, it is expected that seed health quality improves in the next years due to a large governmental investment on seed replacement with certified seed. In any case, our results point out to the weaknesses in the formal seed production system that does not allow farmers to perceive the advantage of using high quality seed. Additionally, these results highlight the importance to fine-tune on-farm seed management practices that promote the increase of seed health quality, and to acquire a better understanding of the seed system and seed sources in Ecuador to support the dissemination of high quality seed to farmers.

### Yield losses caused by seed borne pests

As seen in our results, models estimating yield responses were different between seed sources (Table 4). PVS and PYVV explained yield variation when seed was self-provided by farmers, black scurf when seed was acquired from the market, and PVS and mechanical damage when seed was obtained from the formal system. Additionally, the effect of seed health quality per variety over yield responses was different (Table 4). For instance, the regression model for Chaucha Roja (R^2^ = 9%) identified that the presence of PVS and severity of black scurf in the seed tuber explained yield variation, while for INIAP-Fripapa, only the presence of mechanical damage had an influence (Table 4).

Several studies have shown an increase on yield up to 50% when using seed tuber in optimal physiological and health conditions in comparison with farmer seed tubers (Alvarez 1988; Bertschinger et al. 1995; Wissar 1995; Andrade et al. 2008; Gildemacher et al. 2011; Cesar Garzón 2014). However, these studies have ignored the complexity of interactions (synergy and antagonism) between seed-borne pests and pathogens in multiple contexts (e.g. different soils, environments, and host-pest-pathogen interactions). Inconsistencies of our models (Table 4) suggest that current methodological approaches to understand how seed health quality correlates with yield variation in multiple contexts need still to be further improved, and highlight the relevance of understanding interactions between the different varieties and the seed-borne pest/pathogens to establish indicators that define seed health quality. Similar considerations were mentioned by Thomas-Sharma et al. (2016).

PVS was consistently identified in our models (Table 4) as one the main drivers for yield decline during the period 2010-2011. Additionally, our models per variety estimate that losses caused by this virus range from 27 to 43%. Losses in yield caused by this virus alone are variable but reports mention that it is able to reduce up to 20% of yield (Manzer et al. 1978; Hooker 1981). Special attention need to be given to this virus since the results of our survey revealed a high level of incidence (Table 3 and Table 4).

PYVV was identified as one of the factors explaining a positive effect on yield variation (partial R^2^ of the model for seed self-provided =2%; and partial R^2^ for the model of INIAP-Gabriela = 4%). In our case, this effect was driven by a single variety, INIAP-Gabriela, in which infected plants (see Incidence of PYVV on Table 3) showed a higher yield (2689.2 g plant^-1^; sd = 811.5) than healthy plants (2222.5 g plant^-1^; sd = 696.6) (t-test: p-value < 0.01;

Figure **2**). Salazar (2006) and Guzmán-Barney et al. (2012), on the contrary, indicated that yield is highly reduced in the presence of PYVV and is an emerging disease in Andes. The unusual response of INIAP– Gabriela to PYVV proposes a promising case of investigation to understand this interaction, but also other interactions due to the fact that this variety registered the highest seed-borne pest and pathogen diversity (Figure 1 and Table 3) and the highest yield (Table 4).

**Figure 2.**
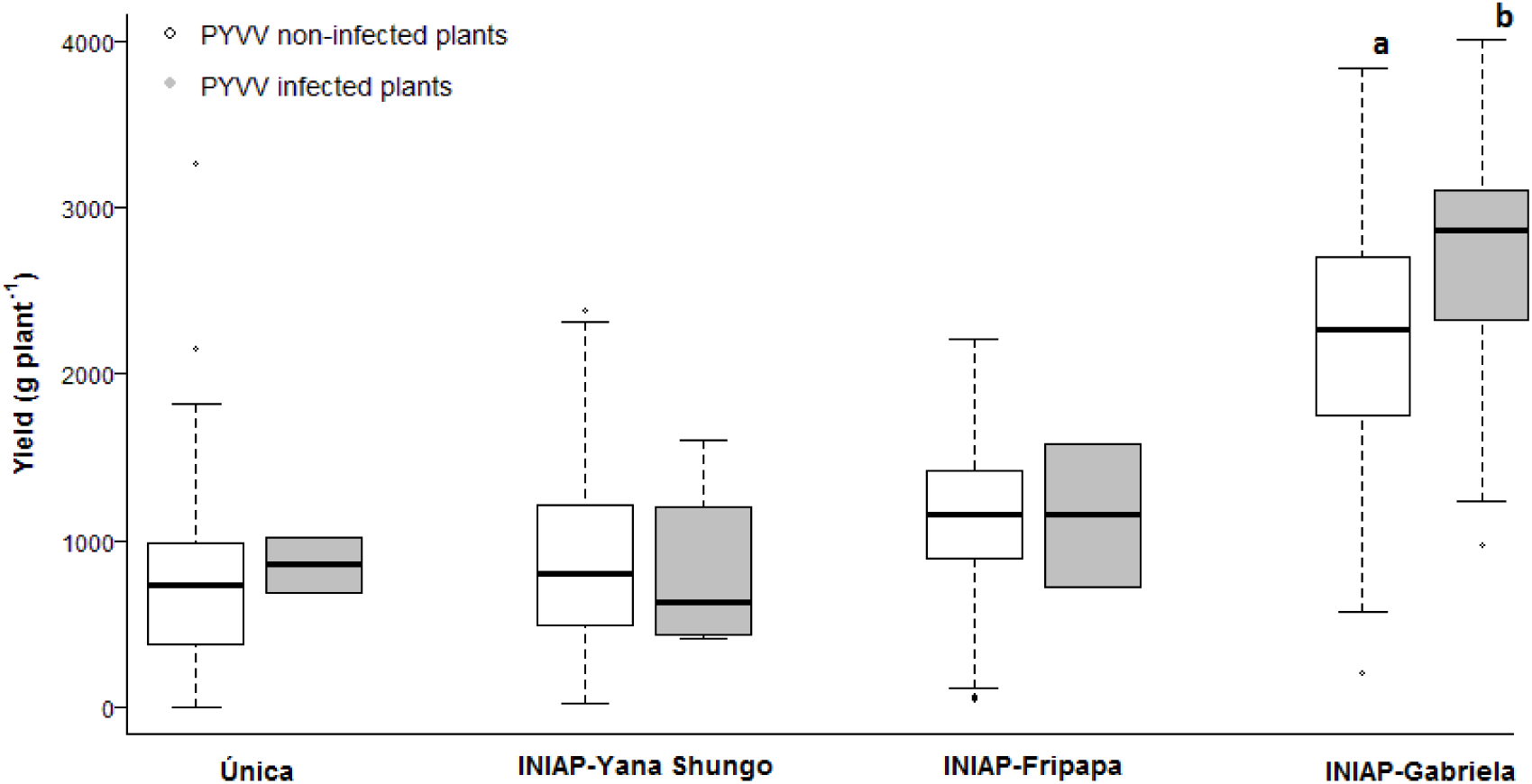
Effect of the presence of PYVV on yield in 4 potato varieties. Yield difference in the variety INIAP-Gabriela was analysed by a t-test. Different letters indicate significant differences according to the t-test (p-value < 0.01). The incidence of PYVV in each variety is in Table 3.

*R. solani* is able to induce yield losses up to 30% on farmer fields (Banville 1989). However, the influence of this pathogen is remarkably affected by the weather and the on-farm management practices (James and McKenzie 1972; Gudmestad et al. 1979; Hill and Anderson 1989; Atkinson et al. 2010). During our experiment, losses in yield caused by *R. solani* were not detected, although, the model detected this pathogen as an important factor driving increase on yield. It is likely that the lack of enough inoculum density induced an increase on yield since the average severity of black scurf on the seed tuber was 3.23% of black scurf (Table 3). Unfortunately, there is no information supporting these findings.

The presence of mechanical damage was relevant in our study (partial R^2^ of the model for seed obtained from the formal seed system =2%; and partial R^2^ for the model of INIAP-Fripapa and INIAP-Yana Shungo = 9% and 11%). The presence of mechanical damage had a negative influence on the yield for the models of the seed obtained from the formal system and INIAP-Fripapa; while it had a positive influence on the yield for the model of INIAP-Yana Shungo. These contradicting results might be due to the fact that mechanical damages were not characterized and only register as either presence or absence. The development of indicators to quantitatively describe mechanical damages could have helped to better estimate losses in yield caused by mechanical damages present on seed tubers.

Despite the fact that our experiment was performed only once without no replicates, our models show that there are several factors affecting yield. We should be cautious at the moment of using this information. Further research is necessary to understand how the seed health quality affects yield taking into account genotype characteristics and different complex conditions (implementing different replicates of the experiment). We believe that resistance to seed borne pathogens should be explore among Ecuadorian varieties to contribute to the improvement of seed health quality in the hands of smallholder farmers.

## Acknowledgements

This research was carried out by the *project “Strengthening systems for native seed potato in Bolivia, Ecuador and Perú”* supported by the McKnight Foundation. We want to acknowledge the support of Fausto Yumisaca, Edwin Pallo, and Fabián Montesdeoca for assistance in the field experiments and fruitful comments; Carlos Barahona and Arturo Taipe for statistical support; and Marcelo Vinueza for technical assistance in the laboratory and glasshouse.

### Author’s contributions

J. Andrade Piedra designed the experiments; I. Navarrete and N. Panchi performed the experiments and analyzed the data; I. Navarrete and J. Andrade-Piedra wrote the article; P. Kromann and G. Forbes provided technical comments and edited the article.

